# Gills are not used for gas exchange in the suspension-feeding hemichordate *Protoglossus graveolens*

**DOI:** 10.1101/2023.08.22.553704

**Authors:** Michael A. Sackville, Christopher B. Cameron, Colin J. Brauner

**Affiliations:** Marine Biological Laboratory, Bay Paul Centre, Woods Hole MA, USA; Département de Sciences Biologiques, Université de Montréal, Montréal PQ, Canada; Department of Zoology, University of British Columbia, Vancouver BC, Canada

## Abstract

The gills are hypothesized to play a key role in early vertebrate evolution by replacing the skin as the primary site of gas exchange. In this scenario, water flow across the gills used for suspension feeding in ancestral deuterostomes was coopted for breathing in stem vertebrates to facilitate the evolution of larger, active fishes. This hypothesis is supported by a stem-vertebrate origin for structures that increase gill capacity for breathing. However, these structures might have instead enhanced an already dominant capacity at the gills of invertebrate deuterostomes rather than mark a shift from the skin. To test this, we characterized gill function for gas exchange in the suspension-feeding hemichordate acorn worm *Protoglossus graveolens*. We measured oxygen uptake and ammonia excretion in whole worms and worm halves with or without gills at 10°C and during an acute thermal challenge at 20°C to maximize gill recruitment. Gills did not enhance oxygen uptake or ammonia excretion, suggesting they are not a primary site of gas exchange. This is the first test of gill function for gas exchange in a suspension-feeding invertebrate deuterostome, and it provides essential support for the long-hypothesized vertebrate origin of breathing at gills and its role in early vertebrate evolution.

## 1. Introduction

The origin and evolution of early vertebrates is characterized by a transition from small, worm-like organisms to larger, more active fishes (Cameron et al. 2000). The gills are hypothesized to play a key role in this iconic transition by replacing the skin as the primary site of breathing (Gans & Northcutt 1983; Northcutt 2005; Brauner & Rombough 2012). In this scenario, water flow across the gills used to filter feed by ancestral deuterostomes was exploited and enhanced for breathing by pharyngeal adaptations in stem vertebrates. This relaxed the worm-like constraints on body form and activity imposed by breathing at the skin, thereby facilitating the evolution of larger and more active fishes.

Support for this scenario is compelling but incomplete. Fossil and developmental studies support a stem-vertebrate origin for structures that increase gill capacity for breathing (Shu et al. 1999; Xian-guang et al. 2002; Mongera et al. 2013; Green & Bronner 2014; Morris & Caron 2014; Green et al. 2015; Gillis & Tidswell 2017). However, it is unknown if these structures mark a true shift from skin to gills for breathing, or if they simply enhance an already dominant capacity at the gills. In vivo measurements in the hemichordate *Saccoglossus kowalevskii* suggest that gills are not the primary site of breathing in this key invertebrate outgroup (Sackville et al. 2022). However, this is the only invertebrate deuterostome in which gill function for gas exchange has been tested. Moreover, aspects of *S. kowalevskii*’s feeding mode suggest that it might be expected to have a lower capacity for gas exchange at gills than other outgroup members.

Indeed, *S. kowalevskii* does not rely on pharyngeal water flow to feed like the proposed vertebrate ancestor and many other invertebrate deuterostomes (Cameron et al. 2000; Cameron 2005; Lowe et al. 2015; Li et al. 2023; Nanglu et al. 2022). As a deposit-feeding filter feeder, *Saccoglossus* instead uses its long proboscis to capture benthic sediment with mucus (Miller 1992; Gonzalez & Cameron 2009). Captured sediment is then moved through the mouth to the pharynx by cilia, where excess water is squeezed from the ingested material and out through the gill slits to produce a concentrated mucus-food cord. This differs from suspension-feeding filter feeders, where a cilia-driven current moves much larger amounts of water into the pharynx and out through the gill slits to filter fine food particles from the water column (Gonzalez & Cameron 2009). Because the deposit-feeding *S. kowalevskii* does not rely on pharyngeal water flow to feed, it may lack the ability to generate sufficient water flow or “ventilation” to breathe with gills relative to suspension feeders. The absence of gas exchange at the gills of *S. kowalevskii* might therefore be a specialized trait linked to feeding mode rather than a general representation of gill function in this outgroup. Robust support for a vertebrate origin of breathing at the gills thus requires further investigation in more invertebrate deuterostomes, and preferably those that suspension feed as well.

Here, we address this knowledge gap by testing gill function for gas exchange in the hemichordate *Protoglossus graveolens. Protoglossus graveolens* closely resembles the morphology and ecology of *S. kowalevskii*, but suspension feeds (Gonzalez & Cameron 2009) and has a body size that is twice as large (wet mass of 0.236 g versus 0.105 g; Sackville et al. 2022). We hypothesize that this suspension feeding machinery and larger body size facilitate and increase gill recruitment for breathing. To test this hypothesis, we leveraged the regenerative capacity of *P. graveolens* to separate worms into viable halves with and without gills for respirometry. Measurements were made at 10°C and during an acute thermal challenge at 20°C to stimulate maximum gill recruitment. We predicted that whole worms and worm halves with gills would have greater rates of oxygen uptake and ammonia excretion than worm halves without gills. This method for isolating gill function was previously validated in *S. kowalevskii* and chosen because acorn worms were too fragile for use in a divided chamber with whole animals (Sackville et al. 2022).

## 2. Methods

### (a) Animal husbandry

*Protoglossus graveolens* were collected from Lowe’s Cove near the Darling Marine Center in Walpole, Maine, USA, and shipped to the University of British Columbia’s Point Grey campus in Vancouver, Canada. Worms were held at 10°C on a 12:12 light:dark photoperiod in static seawater tanks with sandy substrate (10 worms per 40 L tank; 34 ppt artificial seawater from Instant Ocean salt mix, USA). Worms had a mean wet mass of 0.236±0.021 g (n = 27) and were acclimated for 5 days before experimentation.

### (b) 10°C protocol

Eight whole worms were placed in acrylic respirometers (4 ml) in an aerated water bath at 10°C. Each respirometer had a stir bar, overflow tube and needle type oxygen probe (Presens, Germany). Respirometers were left open for 1 h to acclimate worms, then sealed to record the reduction in *P*O_2_ until reaching ∼80% saturation. Worms were then removed from respirometers, cut in half with a transverse section posterior to the gill pores as in Sackville et al. (2022), and moved to 10 ml aerated chambers at 10°C to recover overnight. Once recovered, worm halves were returned to separate respirometers to record *P*O_2_ as before. Following these recordings, worm halves were placed in 20 ml aerated chambers where 1 ml water samples were taken at 10 min and 12 h. Water samples were frozen at -20°C for later measurement of total ammonia as described below. Six additional naive worms were also placed in 20 ml chambers for measurement of ammonia excretion in whole animals. At trial completion, all worms were euthanized with a lethal dose of MS-222 and weighed.

The posterior half of one worm was lost before weighing, which prevented any calculation of *M*O_2_ or *M*NH_3/4_^+^ for it and its corresponding whole worm. All measurements for this worm and its halves were discarded from analyses.

### (c) 20°C protocol

Eight whole worms were warmed from 10°C to 20°C overnight in a water bath at 2.5°C·h^−1^. Worms were then subjected to the same protocol as outlined above for 10°C, but at 20°C. 20°C was chosen as an ecologically relevant thermal challenge that would significantly increase oxygen demand from 10°C, but not result in mortality over 24 h. This temperature was chosen based on prior work with the enteropneust *Saccoglossus kowalevskii* that recommended rearing temperatures not exceed 24°C (Lowe et al. 2004).

### (d) O_2_ uptake

Slopes were taken from the linear portions of traces recorded for each respirometer with OXY-4V2_11TX data acquisition software (Presens, Germany). Slopes were adjusted with background respiration values from blank trials to yield a Δ*P*O_2_·h^−1^ (mm Hg O_2_·h^−1^). These values were used with respirometer volume, worm wet mass and O_2_ solubility values (μmols O_2_·ml^−1^·mm Hg^−1^ for respective temperatures from Boutilier et al. 1984) to calculate oxygen uptake rates (*M*O_2_; μmols O2·g^−1^·h^−1^).

### (e) NH_3/4_ ^+^ excretion

Water samples were analysed for total ammonia (NH_3/4_ ^+^) colourometrically (Verdouw et al.) in triplicate using a SpectraMax 190 microplate reader (Molecular Devices, USA) as in Sackville et al. (2022). The difference in ammonia concentration over trial duration was used to calculate rate of change in ammonia concentration (nmols NH_3/4_^+^·ml^−1^·h^−1^). This rate was used with respirometer volume and worm wet mass to calculate ammonia excretion rates (*M*NH_3/4_^+^; μmols NH_3/4_ ^+^·g^−1^·h^−1^).

### (f) Statistics and reproducibility

Data were analysed with Prism 9 for macOS (Version 9.3.1; GraphPad Software Inc., USA). Gas exchange in whole and fragmented worms was analysed with one-way ANOVA and Tukey’s post-hoc test. All data passed tests of normality and equal variance, but some required transformation (1/x for *M*O_2_ & *M*NH_3/4_ ^+^ at 10°C; log(x) for *M*NH_3/4_ ^+^ at 20°C).

Sample sizes were determined from prior studies where similar methods found meaningful biological effects. Animals were randomly assigned to experimental treatments, but blinding was impossible as water temperature required monitoring and manipulation during experimentation. All other data collection and analyses were blind to treatments.

## 3. Results & discussion

Contrary to our hypothesis, our results indicate that gills are not a primary site of gas exchange in the suspension-feeding hemichordate *Protoglossus graveolens*. Mass-specific rates of oxygen uptake (*M*O_2_) and ammonia excretion (*M*NH_3/4_^+^) for worm halves without gills were at least as high as rates for whole worms and worm halves with gills (Fig. 1a, b). This was true at both temperatures despite a 1.9- and 2.6-fold increase at 20°C for *M*O_2_ and *M*NH_3/4_^+^, respectively. Rates were similar to those reported for the deposit-feeding hemichordate *Saccoglossus kowalevskii*, suggesting that our experimental preparation did not affect worms in any unexpected way (Sackville et al. 2022). Furthermore, temperature effects (*Q*_10_; ratio between rates over a span of 10°C) lower than 2 for *M*O_2_ and above 2.5 for *M*NH_3/4_^+^ support 20°C as a suitable challenge near an upper thermal limit (Schulte 2015).

**Figure 1:**
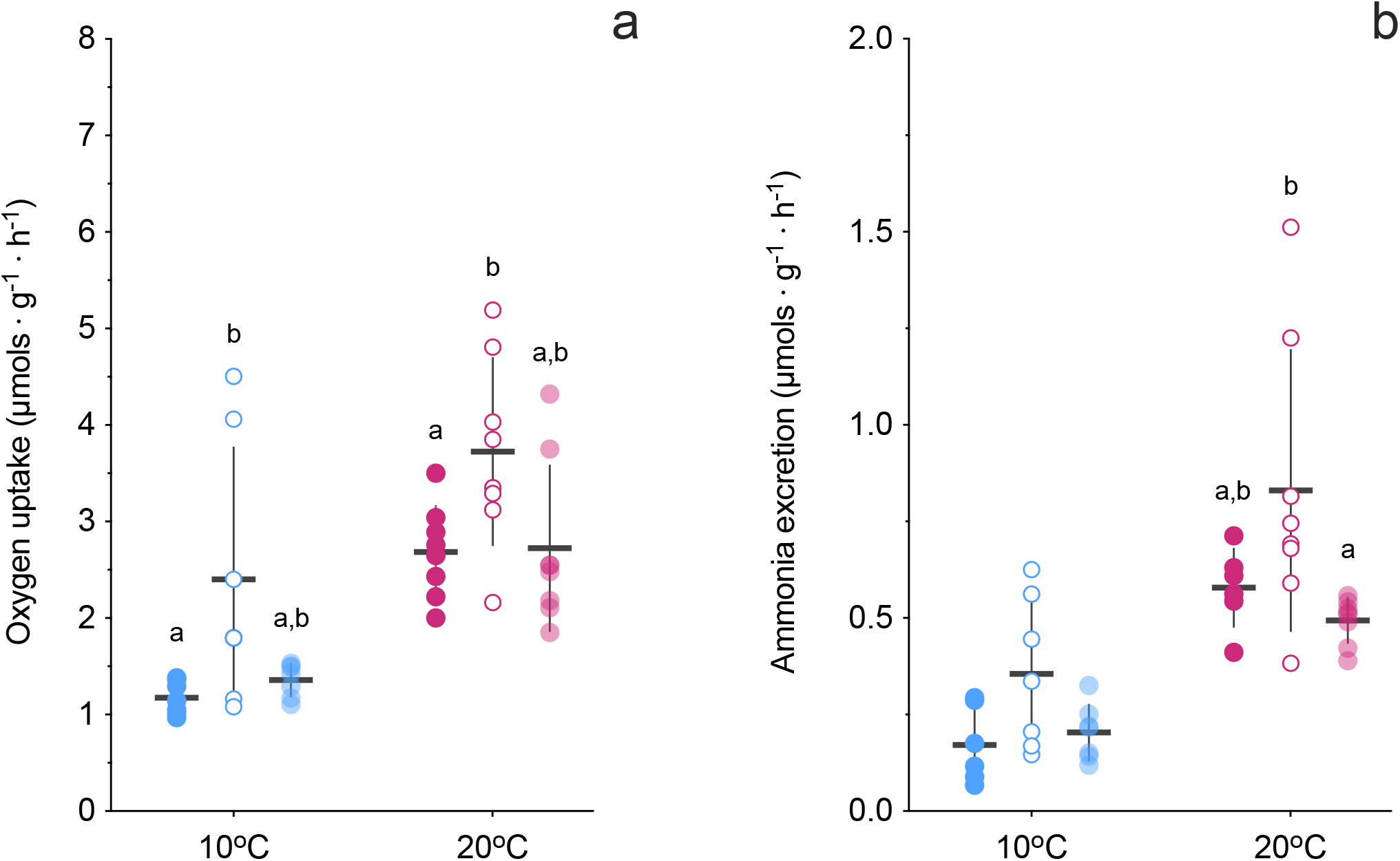
Mass-specific rates of (a) oxygen uptake and (b) ammonia excretion in whole worms (solid symbols), worm halves without gills (open symbols) and worm halves with gills (shaded symbols) at 10°C (blue) and 20°C (red). Data presented as means +/− sd with individuals superimposed. Letters that differ indicate significant differences within treatments (one-way ANOVA with Tukey’s test, *P*<0.05, n=6-8).

In some treatments, *M*O_2_ and *M*NH_3/4_ ^+^ were higher in worm halves without gills than whole worms or worm halves with gills (Fig. 1; *M*O_2_ at 10°C and 20°C, *M*NH_3/4_ ^+^ at 20°C). However, this appears to be associated with variation in worm fragment mass (Fig. 2). Indeed, oxygen uptake scales with fragment mass to the power of ∼3/4 at both temperatures (Fig. 2a), and ammonia excretion to the power of ∼2/3 at 20°C (Fig. 2b). It is unclear if the changes to *M*O_2_ and *M*NH_3/4_ ^+^ following worm fragmentation are caused by simple changes in surface area to volume ratio or other factors, and this is beyond the scope of our work. However, these results suggest that acorn worms and their regenerative capacity could be a powerful model system in which to test the mechanisms that underlie metabolic scaling, a topic of enduring controversy across biological disciplines (Rubner 1883; Kleiber 1932; Schmidt-Nielsen 1984; West et al. 1997; White & Seymour 2003; Savage et al. 2004; Glazier 2010; White & Kearney 2013; White et al. 2022; Glazier 2022; Freose & Pauly 2023).

**Figure 2:**
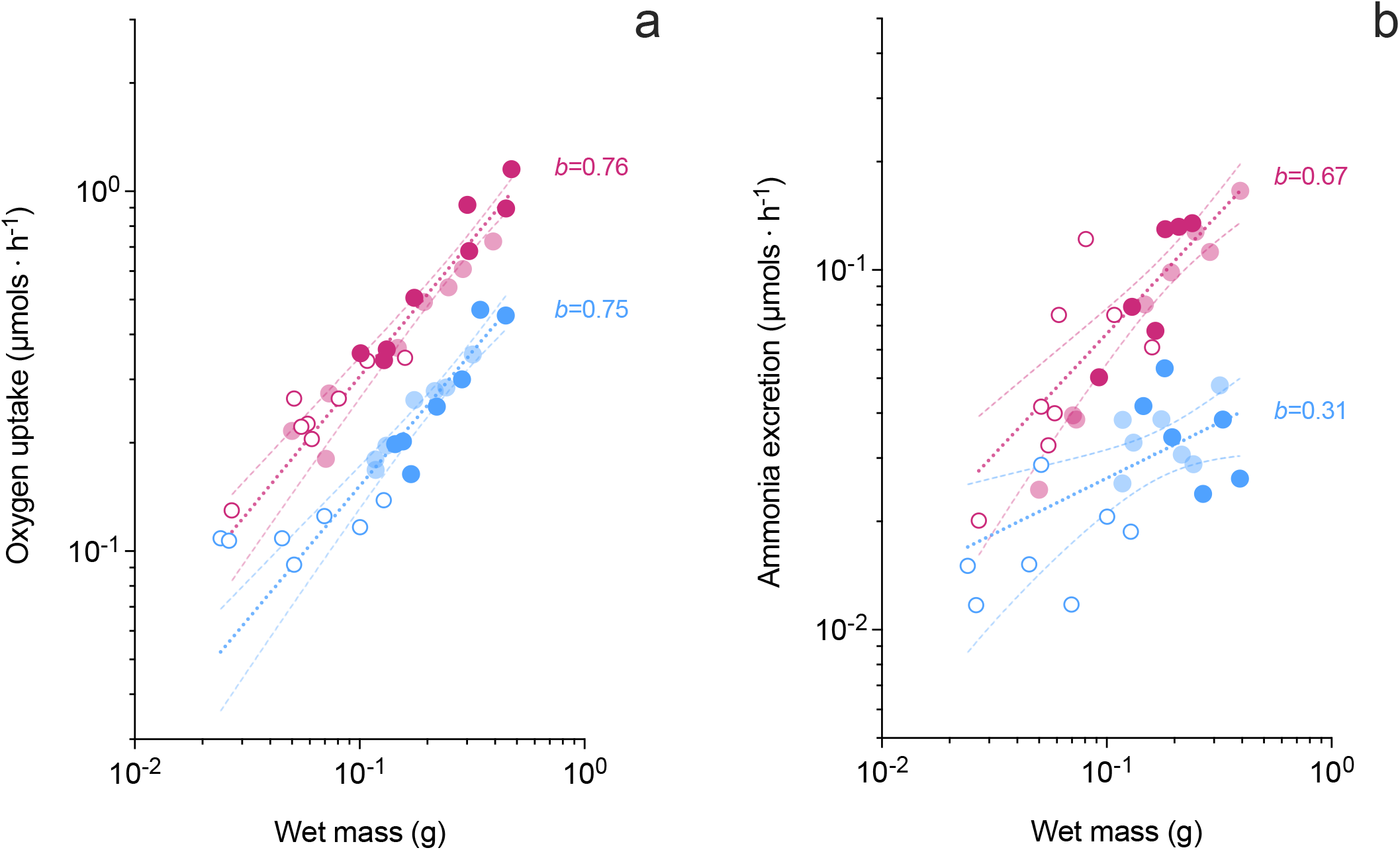
Rates of (a) oxygen uptake and (b) ammonia excretion expressed as a function of wet mass in whole worms (solid symbols), worm halves without gills (open symbols) and worm halves with gills (shaded symbols) at 10°C (blue) and 20°C (red). Allometric curves +/-95% C.I. are fitted to data for each temperature. Oxygen uptake at 10°C (*b*=0.75, r^2^=0.90, n=21), 20°C (*b*=0.76, r^2^=0.92, n=24); ammonia excretion at 10°C (*b*=0.31, r^2^=0.36, n=20), 20°C (*b*=0.67, r^2^=0.74, n=22).

This is the first test of gill function for gas exchange in a suspension-feeding invertebrate deuterostome, and only the second invertebrate deuterostome to be tested at all. Together with prior findings in *S. kowalevskii* (Sackville et al. 2022), these results suggest that the gills of hemichordate acorn worms do not play a significant role in gas exchange whether they suspension feed or deposit feed. This is essential and compelling support for a vertebrate origin of gas exchange at gills, as these data are the only functional measurements from this key invertebrate outgroup. However, functional measurements are still missing from other key outgroups within the deuterostomes that possess pharyngeal gills, including the cephalochordates and urochordates. These taxa are more closely related to vertebrates than hemichordates and should be targeted for future study. Functional measurements that demonstrate an absence of gas exchange at the gills of these taxa will allow us to more confidently resolve the origin of breathing at gills to the vertebrate stem, and to determine if the pattern observed in hemichordate acorn worms truly represents a shared ancestral condition among all invertebrate deuterostomes.

## Contributions

MAS, CJB and CBC conceived the study. MAS performed all experiments and analyses and wrote the manuscript. All authors provided manuscript edits and approved the final version.

## Ethics

All animal care and experimentation conformed to the guidelines set by the Canadian Council on Animal Care (CCAC) and was approved by the University of British Columbia’s Animal Care Committee (ACC) under the animal use protocol A19-0284.

## Data accessibility

All data are provided in the supplemental material.

